# Sfaira accelerates data and model reuse in single cell genomics

**DOI:** 10.1101/2020.12.16.419036

**Authors:** David S. Fischer, Leander Dony, Martin König, Abdul Moeed, Luke Zappia, Sophie Tritschler, Olle Holmberg, Hananeh Aliee, Fabian J. Theis

## Abstract

Exploratory analysis of single-cell RNA-seq data sets is currently based on statistical and machine learning models that are adapted to each new data set from scratch. A typical analysis workflow includes a choice of dimensionality reduction, selection of clustering parameters, and mapping of prior annotation. These steps typically require several iterations and can take up significant time in many single-cell RNA-seq projects. Here, we introduce sfaira, which is a single-cell data and model zoo which houses data sets as well as pre-trained models. The data zoo is designed to facilitate the fast and easy contribution of data sets, interfacing to a large community of data providers. Sfaira currently includes 233 data sets across 45 organs and 3.1 million cells in both human and mouse. Using these data sets we have trained eight different example model classes, such as autoencoders and logistic cell type predictors: The infrastructure of sfaira is model agnostic and allows training und usage of many previously published models. Sfaira directly aids in exploratory data analysis by replacing embedding and cell type annotation workflows with end-to-end pre-trained parametric models. As further example use cases for sfaira, we demonstrate the extraction of gene-centric data statistics across many tissues, improved usage of cell type labels at different levels of coarseness, and an application for learning interpretable models through data regularization on extremely diverse data sets.

## Background

Many single-cell data sets are currently published in different databases in different formats, such as custom formats on GEO, manuscript supplements with tables of cell type annotations, or streamlined formats on Human Cell Atlas servers. Similarly, many parametric models for data integration, cell type annotation and other tasks are published with their own user interface. The lack of streamlined data and model access inhibits data and model re-use and makes comparative analyses and benchmarks work intensive. We identify two core issues with the current state of data and model re-use in single-cell genomics. Firstly, in smaller data sets, rare cell states can often only be properly analysed after integration with larger reference atlas data sets. This integration is time intensive and requires a prior analysis of the reference data set. The effectiveness of this approach depends on the reference atlas chosen. With a growing number of available reference data sets^1,2^, the choice of integration method and reference data set become increasingly hard to explore for analysts^3^. Secondly, data processing and cell type annotation are repeated elements of these pipelines that are time intensive for analysts because of the complexity of the pipelines used^4^. Both computing an embedding and clustering require basic preprocessing, such as scaling, log-transformation and highly variable feature selection. This data processing, also called feature engineering, is typically necessary both for basic embeddings such as t-SNE or uniform manifold approximation and projection^4^ (UMAP) but also for embeddings from autoencoders such as DCA^5^ and scVI^6^. Moreover, cell type annotation requires a high level of domain expertise as annotation resolution depends on quality of the data and project requirements and, because cell type ontologies are currently under development and therefore may change over the time scale of a typical analysis project.

We argue that zoos of pre-trained models can alleviate these problems by replacing processing steps that are usually manually tuned by analysts with standardized parametric models that correspond to entire processing pipelines. First, similar models can be trained on different data sets or collections, allowing analysts to navigate different reference data sets easily. Second, a zoo eases model sharing through a unified front-end. The idea of model sharing has been successfully applied in other fields including natural language processing and computer vision and in geomics with kipoi^7^ for sequence-based models. Here, we introduce sfaira, a versatile repository that serves pre-trained scRNA-seq models. To train these models across tissues and organisms, we coupled the zoo with a data repository that includes data sets from multiple data providers with unified annotations. This data and model zoo permits streamlined access to data sets and pre-trained models. The presented sfaira framework defines a common nomenclature that covers feature spaces, data sets, and cell type ontologies. We leverage this data zoo to train models in an automated fashion across large numbers of tissues in two species and propose a mechanism that automatically accounts for different cell type annotation resolutions in cell type classifier models. Current model zoos are model class centric, thereby impeding side-by-side usage of different models, such as different autoencoder topologies. The sfaira model zoo is designed to be model agnostic and to simply be as a unified front-end for serving and receiving models, thereby enabling transfer of models from developers to users.

In addition to these practical advantages of a data and model zoo, we also address the issue of interpretability and generalizability of models. We provide a size factor-normalised, but otherwise non-processed feature space, for models so that all genes can contribute to embeddings and classification and the contribution of all genes can be dissected without the issue of removing low variance features. We show that this approach allows us to relate the dimensions of the latent space to all genes. We also present models that have been fitted without covariates, such as organ or experimental assay, on extremely diverse data sets. We argue that such models require higher abstraction on the gene space compared to models that use covariates to remove variation between data sets. To our knowledge, this is the first instance in which such models could successfully be trained. Altogether, we expect sfaira to provide the important service of model reuse and broad model profiling across a diverse range of unified data sets.

## Results

### Sfaira provides data sets, models, annotation and model parameters within a unified framework

Sfaira provides data, model, and parameter estimate access as a data and model zoo (Fig. 1a). Firstly, the data zoo contains data set-specific loader classes to query data from the actual diverse data providers, which mirror data reading scripts and make these scripts sharable and reproducible. This data zoo is scalable because data loaders can be easily shared. Currently, as of December 2020, sfaira encompasses 38 studies with 233 data sets and 3.1 million cells (Fig. 1b). This data loader implementation allows streamlined querying of data sets based on meta features, such as organism or tissue sampled and experimental protocol. We couple this data representation to a cell type ontology, such that cell type annotation between data sets can be easily related to each other. The gene space is explicitly coupled to a genome assembly to allow controlled feature space mapping.

**Figure 1:**
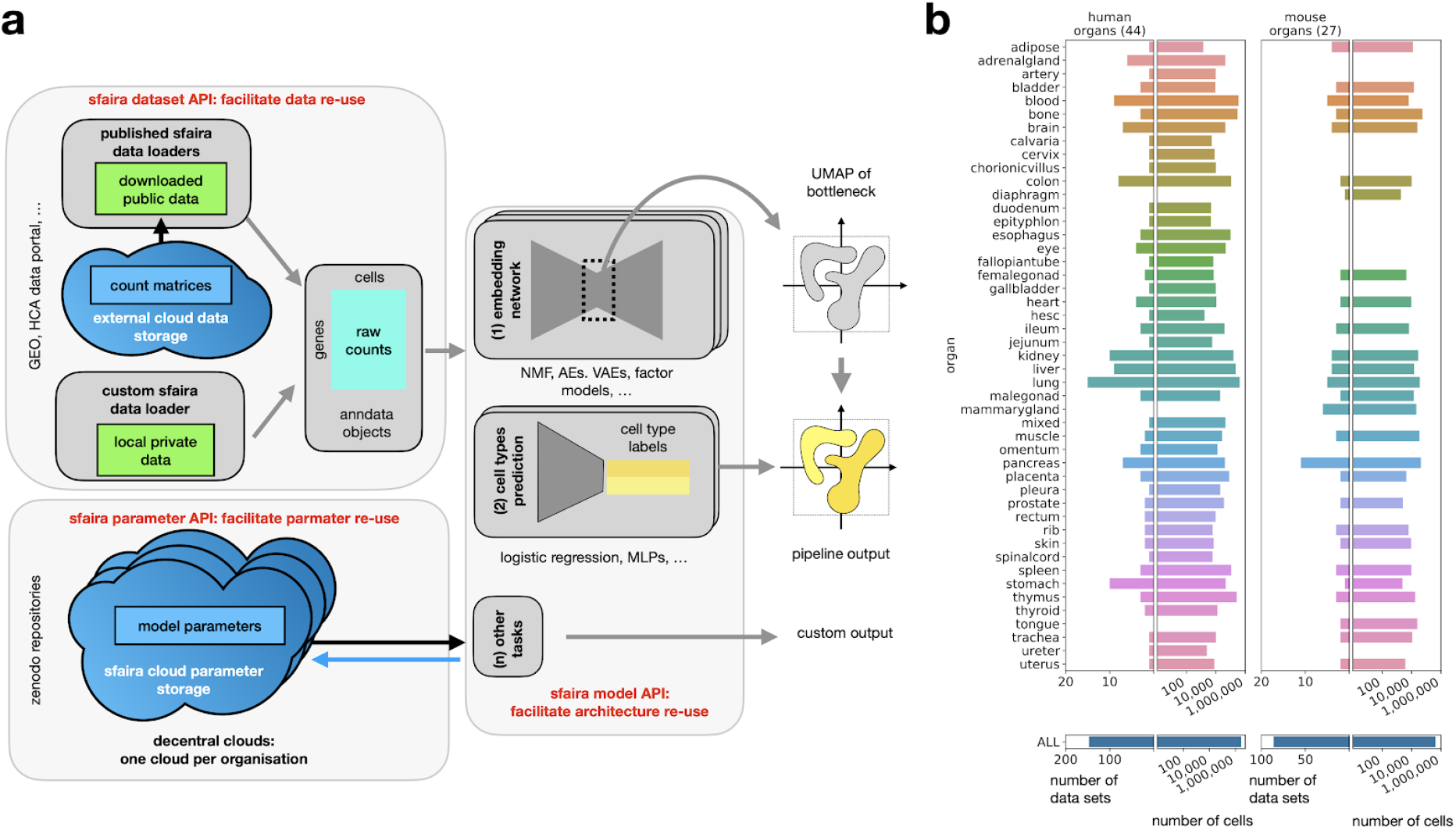
Sfaira is a data and model zoo that automates common steps in exploratory single-cell RNA-seq analysis. **(a)** Overview workflow of sfaira data and model API. Data set files are currently stored in cloud databases that can be interfaced with the sfaira data API to give streamlined AnnData objects that can be used for analysis or model fitting. The sfaira model API can consume these data sets to produce automatised analyses by querying parameter estimates from pre-trained models, stored on cloud servers via the sfaira parameter API. Example analysis steps that are automatised are embedding computation and cell type annotation. **(b)** Summary of the current state of sfaira data zoo for mouse and human single-cell RNA-seq samples, representing 233 data sets across 45 organs and 3.1 million cells in two organisms.

Secondly, the model zoo part of sfaira consists of a unified user interface and model implementations, not requiring the user to understand technical differences between models such as supervised cell type prediction models, matrix factorisations, and variational autoencoders. It is often desirable to use pre-trained models during analysis. For this purpose, we couple the pre-implemented models to parameter estimates stored locally or in a cloud database. These parameterizations can easily be queried from within Python workflows and allow streamlined execution of previously published models. The parameter query depends on a global model and parameter versioning system that we introduce with sfaira.

### A scalable data zoo for fast and comprehensive data access across numerous repositories

Important technical challenges with data zoos are reading and performing interactive workflows on large, heterogeneous data set collections that do not fit into memory. We address data loading by using streamlined data-set-specific loader classes that contain data-loading scripts. These classes can be written, maintained, and used in the context of the complex functionalities of parent classes, as well as shared through a single public code base. Moreover, we extend these data-set-specific loader classes to data collection loaders that serve already streamlined data sets. With these, sfaira serves a maximum number of data sets with minimal code effort and high potential for sharing and contributing. Importantly, sfaira only requires a constant amount of code to load data sets, independent of the set of selected data sets. We also introduce lazy loaded representations of data sets that allow users to subset large data set collections before loading desired subsets into memory. Here, we provide functionalities to write larger joint data sets with a streamlined feature space and cell type annotation, which can be handled on disk through AnnData’s disk-backed formats^8^. Last, we also aid data zoo exploration through a web front end that contains a searchable summary of all data sets in the data zoo database (*Availability of data and materials)*.

### Scalable access to data with unified annotations allows for queries of gene and data statistics

Streamlined access to unified, large, cross-study data sets as provided by sfaira allows for easy data statistics queries that can be helpful for putting observations in the context of other data sets. This includes queries for the expression of specific genes or consistent gene interactions across tissues or atlas-scale queries for cell type frequencies. For example, it is often useful to have a reference range for the expression activity of a gene observed with scRNA-seq. This can be done with sfaira via a single straight-forward query, as showcased for *Ins1* scaled expression across organs and cell types in mouse (Fig. 2a). With a few commands, one can establish an active range of expression of this gene in the entire mouse data zoo with 895 cell type observations across 86 data sets. Note that cell type-wise summary metrics are often much more useful for such workflows than cross-data set averages, which are skewed toward frequent cell types and are more useful than extrema, such as maximum expression, which are heavily influenced by the variance of the expression distribution. Cell type-wise metrics can only be easily obtained in a framework that contains a unified cell ontology that is generalized across studies, such as implemented in sfaira. This analysis establishes an active range between 0 and 2,500 counts per 10,000 unique molecular identifiers as an active range for *Ins1* expression in mice, with all expressing cell types located in pancreas data sets.

**Figure 2:**
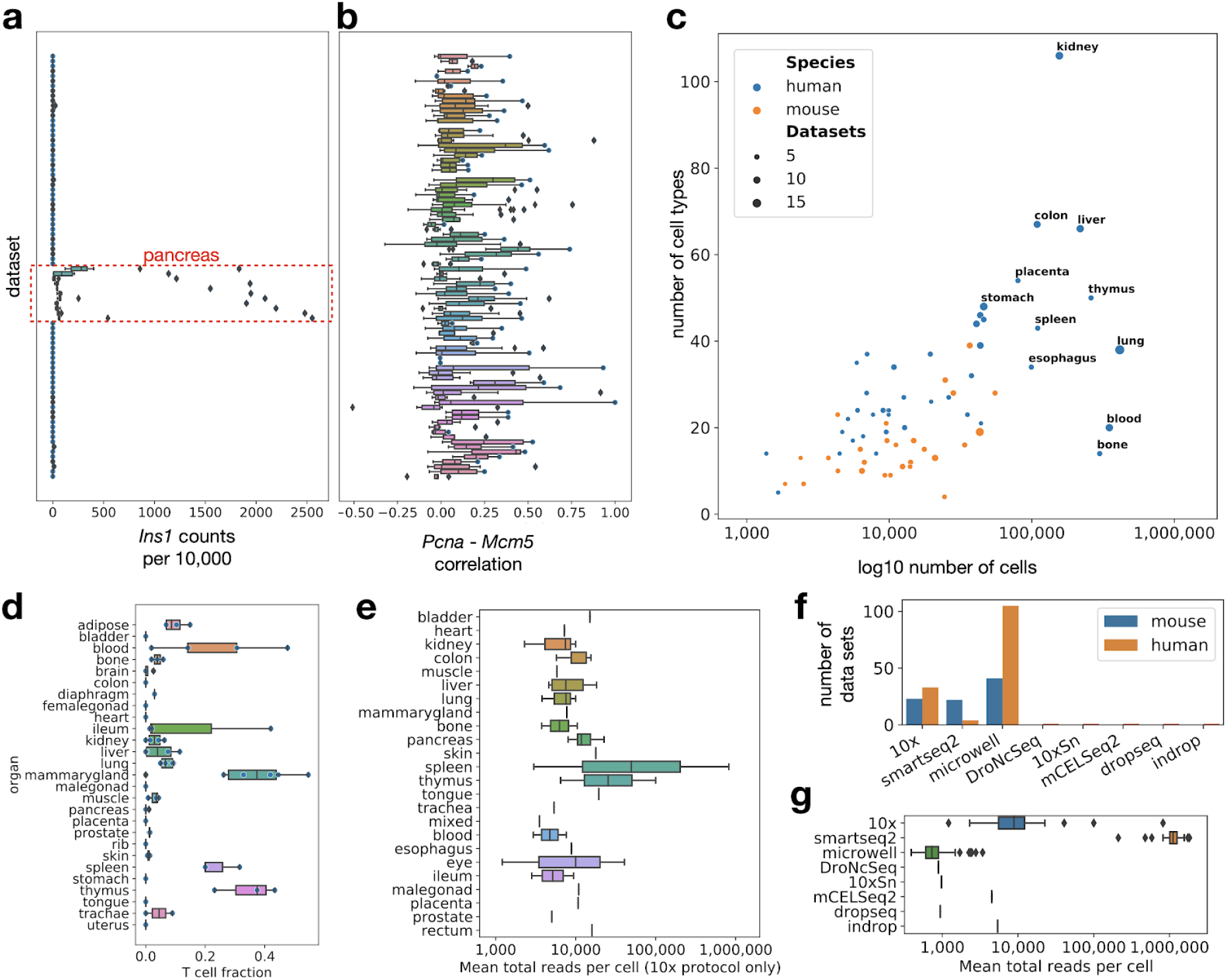
The sfaira data zoo contextualizes data statistics. **(a)** Expression range of an example gene across data sets. Mean normalized expression of *Ins1* by organ and cell type. **(b)** Pearson correlation coefficient between the two cell-cycle-associated genes *Pcna* and *Mcm5* across data sets. Shown is the a boxplot of the distribution over the correlation coefficients for each data set computed per organ and cell type. **(c)** Data sets vary strongly in complexity. Shown is the number of cells versus the number of cell types in the data zoo by organ for both mice and humans. **(d)** Sfaira allows query of cell type fractions in tissue across organisms. Shown is the fraction of T cells per dataset in mice, ordered by organ **(e)** Mean total counts per cell in mouse and human organs for 10x protocol data sets only. **(f)** Number of data sets per experimental protocol. **(g)** Mean number of counts (unique molecular identifiers if available, otherwise reads) per data sets by experimental protocol.

We can also consider gene-gene dependencies. Often, one is interested in the correlation of expression between genes to establish regulatory relationships. As an example, we investigated the correlation of two cell-cycle-associated genes, *Mcm5* and *Pcna* (Fig. 2b), which provides a range for their correlation and an estimate of how often these genes are correlated across tissues. This analysis shows a range of correlation coefficients including many high values above 0.5, which establishes the expression of these two genes are correlated in many cell types, which is to be expected, because the cell cycle is a common biological confounder of transcriptomic states.

As well as these local data characteristics, one can also use a data zoo to obtain global, atlas-level statistics. We consider the total number of cells versus the number of most fine grained cell type labels per organ (Fig. 2c). Such complexity plots could be used to prioritize organs for further cell type discovery. The examples shown here reflect cases in which a streamlined data pipeline makes analyses possible that would otherwise be too time consuming to be conducted on a regular basis. Moreover, one can query the fraction of cells of a particular type, such as T cells, across organs easily (Fig. 2d). We also provided a summary of total reads per cell summary statistics and protocol summary statistics (Fig. 2e-g).

### Sfaira enables automated single-cell data analysis

A core advantage of end-to-end parametric approaches is that they can alleviate the need for feature engineering. This has been a key advancement in image-based deep learning for example^9^. In single-cell analyses, feature engineering describes the early analysis steps starting from count matrices, including normalization, log-transformation, gene filtering, selecting components from principal component analysis (PCA), and batch correction^4,10^ (Fig. 3a). These steps are usually necessary to obtain useful embeddings and clusterings^4^ but are a bottleneck in analysis workflows. Pre-trained embedding models can be used to generate latent spaces that can be used for downstream tasks without prior feature engineering. As an example case, we processed human peripheral blood mononuclear cells (PBMC) data in a standard preprocessing workflow^4,8^ and compared this to a UMAP of a linear model embedding. Both the manual and the learned embedding separated annotated cell types into distinct clusters, which demonstrates that both captured the biological heterogeneity of the system (Fig. 3b). One could judge the learned embedding also based on the reconstruction error of its encoder-decoder model: Here, the linear model achieved a mean negative log likelihood in reconstruction of 0.14. These quantitative metrics on embedding models become important when multiple models are compared. Second, we used automated cell type annotation to label cells to explore whether we could seed data interpretation with a first proposal for cell types. Our cell type predictions from a logistic regression model trained on different data sets identified the same coarse cell types: T cells, antigen-presenting cells, monocytes, and dendritic cells (Fig. 3b). Note that with further additions to the data zoo and improved classifier models trained on these large data sets, these coarse initial annotations will become increasingly fine-grained. This example shows that the combination of pre-trained embedding and cell type classifier models can be used to perform an automated initial analysis of single-cell data, which can then be extended by further in depth analysis according to the scenario. Importantly, this analysis is entirely automated through a few lines of code, thereby reducing the time required to explore the data by a few hours or even days, depending on the expertise and domain knowledge of the analyst. Below, we discuss pre-training details of such cell type classifiers and embedding models that allow these workflows on a large-scale.

**Figure 3:**
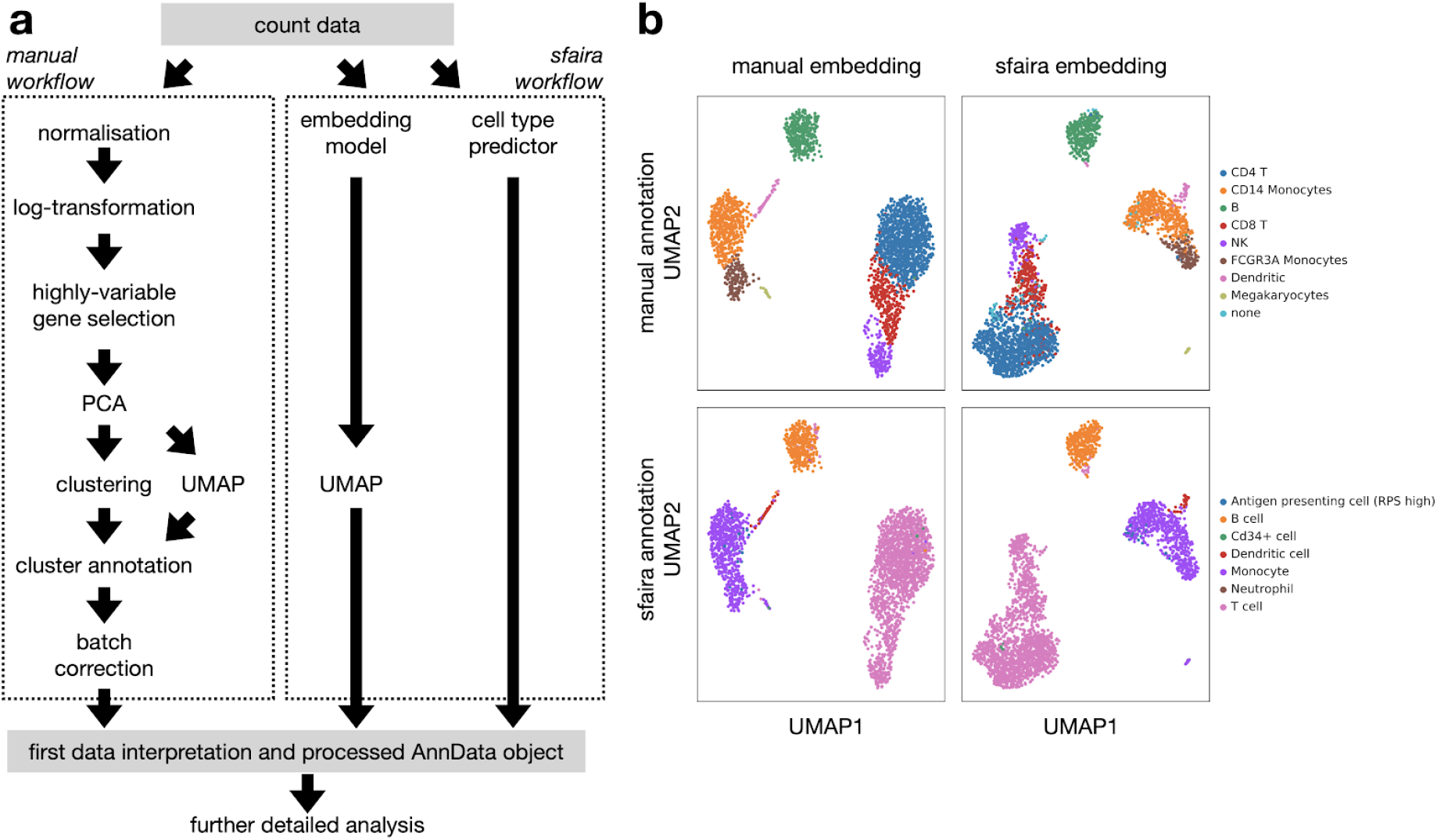
Sfaira automatizes exploratory analysis of single-cell data. **(a)** Manual single-cell data analysis pipeline and automated sfaira pipeline. **(b)** Comparison of manual feature engineering workflow with automated embedding and cell type annotation from sfaira on human PBMC dataset. Shown are UMAPs based on a PCA of an engineered feature space and of the out-of-the-box latent space from a linear sfaira embedding model. Superimposed are cell types previously annotated for this data set and sfaira cell type predictions.

### Sfaira versions decentralized parametric models to allow reproducible model sharing and application to private data

Sfaira currently implements two types of models: (i) unsupervised encoding models that learn a latent representation of data that can be used for visualization, and (ii) supervised models that predict cell type labels (Fig. 1a). However, sfaira’s architecture is sufficiently flexible to integrate other model types that serve additional purposes. For each embedding model, the following metadata is stored. The input feature space, which is defined by the genes in a genome annotation (Fig. 1a), the model topology, the biological specimen (e.g., tissue and species) it was trained for, and an identifier for the training strategy (training data sets and optimization hyperparameters). Cell type classifiers include the above as well as a defined cell type ontology^11^. We broadly categorize embedding model topologies according to popular approaches, such as matrix factorizations, autoencoders, and variational autoencoders. Similarly, we include logistic regression and densely connected neural networks for cell type classification. Each of these can vary in multiple hyperparameters, such as depth, embedding width, activation functions, regularization, and training mode. For a single term in this nomenclature, one may still obtain different models that vary in their parameter estimates, thus requiring a version identifier for the parameter estimate. Taken together, this short model ontology allows for a unique query of a particular model, thereby guiding the usage of such models in a reproducible manner within a model zoo. “celltype_mouse_bladder_mlp_theislab_0.1.1_0.2” would be a celltype classifier model trained on mouse bladder data sets, a multilayer perceptron architecture with hyperparameter version 0.1.1, with parameter version 0.2 and provided by the organization theislab.

We provide an infrastructure for multiple organizations to maintain their own public and private repositories of model weights (model zoos) on servers such as Zenodo or in local directories. These parameter set versions are identified by the organization that performed model training as well as the training data and optimization hyperparameters that this organisation used to train this model. Often, this would result in organizations providing an initial estimate that becomes incrementally updated as new data becomes available or when improved estimates become available in an ongoing grid search across optimization hyper-parameters. Sfaira allows end-users to easily switch between different model types from different model providers, accelerating and democratizing model distribution and access. This reduction in the effort required to quickly implement and compare models will improve decisions on pre-trained model usage. In addition, the decentralized storage of model weights allows this model zoo to quickly react to new developments in the community.

### Generalized cell type prediction within an ontology adjusts for annotation coarseness

A core difficulty for deploying predictive models for cell type labels based on single-cell RNA-seq is that the changing universe of cell types is redefined and new types discovered as part of cellular atlas efforts^12^. We address this issue by defining versioned cell type universes, which are sets of leave nodes of a cell ontology that correspond to the most fine grained cell types that are characterized within a given tissue. The universes are integrated in a cell ontology for each organism (Fig. 1b) and are part of the cell type predictor model version. A second challenge is that cell type annotation from previous studies is often presented at different resolutions. One study might report “leukocytes” in a given tissue while a different study differentiates between “T cells” and “macrophages”. A scalable training framework for cell type classifiers needs to be able to make use of both levels of granularity, as manual re-annotation is extremely time-consuming and may not always achieve the required resolution, depending on data quality. This notion of coarseness relates to the directed acyclic graphs that are typically employed in cell type ontologies. Accordingly, we propose the usage of a variant of cross-entropy loss and an accuracy metric that can dynamically assign observations to single labels or to sets of labels during training and testing (aggregated cross entropy, Fig. 4a, see Methods). Using this approach, we were able to pool cell type annotations from more than 200 public data sets and train predictive models for 27 tissues from mice and 44 from humans across 3.1 million cells at once.

**Figure 4:**
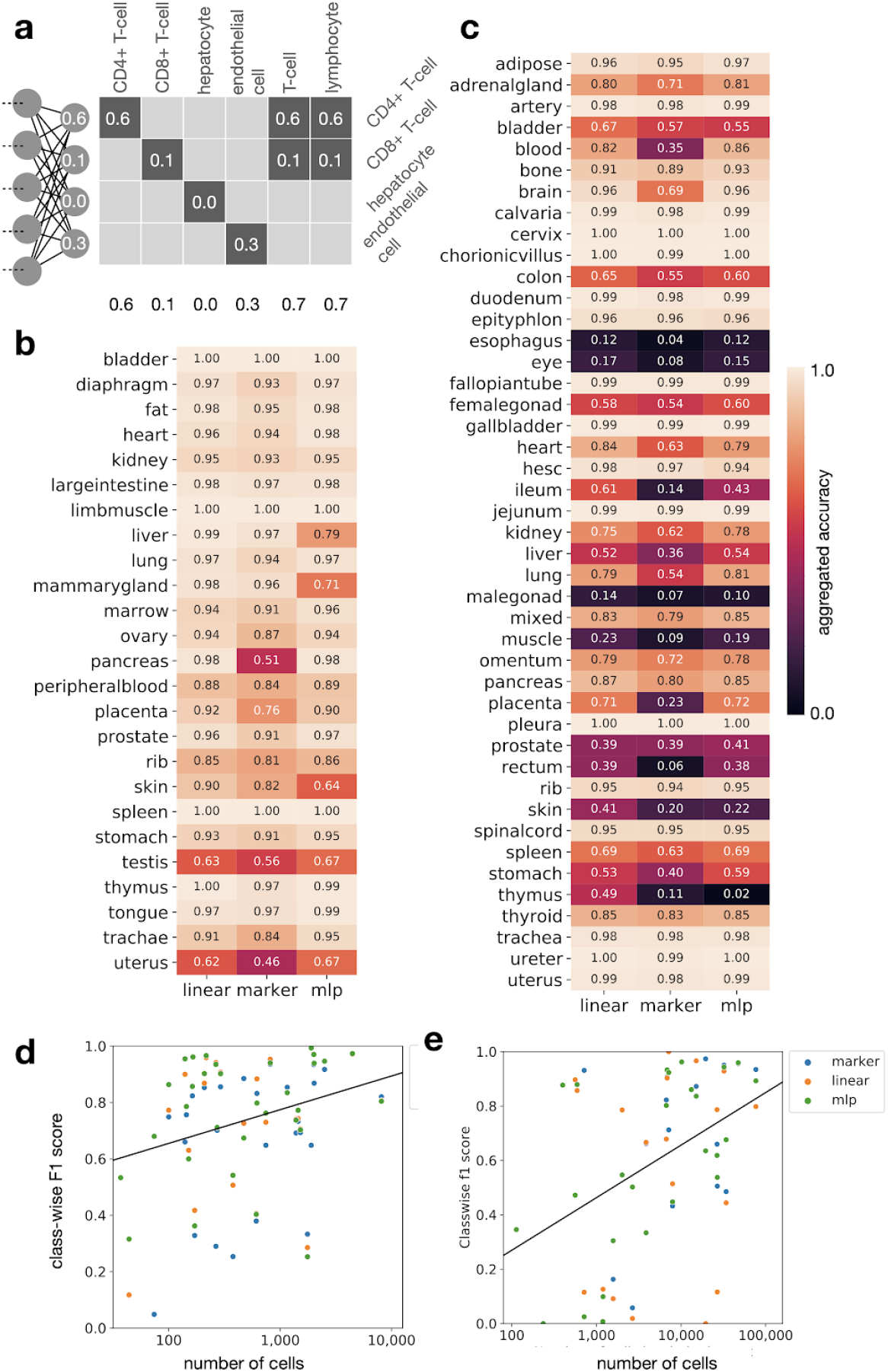
Sfaira allows fitting of cell type classifiers for data sets with different levels of annotation granularity by using cell type ontologies. **a)** Aggregated accuracy and cross-entropy allow for fitting cell type classification models on data sets with heterogeneous annotation coarsity using cell type relations from ontologies (see **Methods**). **(b,c)** Accuracy of cell type classifiers on mouse (**b**) and human (**c**) organs on entirely held-out test-data sets. Linear: Linear classifier (logistic regression), marker: Marker gene-based classifier, MLP: multilayer dense neural network. **(d,e)** Class-wise prediction accuracy correlates with the number of cells in class. Shown are cell type class-wise F1 scores by the number of cell types in that class of cell type classifiers by model on lung data from mice (**d**) and humans (**e**).

It has recently been proposed that cell type prediction can often be performed with linear models^13^. We trained three types of models: logistic regression models, three-layer densely connected feed forward neural networks and a new marker gene-centric linear model (Methods). The newly proposed gene-centric model operates in a learned marker gene space in which each gene is first transformed into a binary on-off state with a sigmoid mapping. Such models are not only easy to interpret, as marker genes contribute equally to the prediction, but they also allow integration of prior knowledge on marker genes via priors for the parameters of the marker state embedding layer. We also argue that the marker-gene-based prediction is more similar to the training data generation (cell type annotation) process and will, therefore, generalize more stably. All models performed well as expected based on previous findings on selected organs (Fig. 4b,c), with a median accuracy of 0.82 in human samples and 0.95 in mouse samples. Our data zoo facilitates training and deployment of these models in a streamlined fashion, thus making cell type predictors easily accessible for all sampled organs and organisms. Using the data zoo, we can easily relate classifier performance to class frequencies (Fig. 4d,e) and can consider individual classes in more detail (Supp. Fig. 12).

### Sfaira serves embeddings from different models

Embedding models compress data to a low-dimensional representation and are at the core of unsupervised data analysis. Usually, data set compressions are visually inspected in 2D dimension reductions, such as from t-SNE or UMAP. Encoder-decoder neural network topologies (autoencoders) extend PCA to non-linear models for this purpose^5,14^. Members of this model class that have been used frequently in the past for representation learning on single-cell RNA-seq are PCA, non-negative matrix factorization^15,16^, autoencoders^5^, and variational autoencoders^6^. Embedding models have been successfully used in the context of transfer-learning^15,16^, a process during which public data are leveraged to improve learned representations. Workflows that use such encoder-decoder models in unsupervised scRNA-seq data analysis usually rely on refitting of the model on each new data set for two core reasons. Useful pre-trained models are difficult to identify in the literature and are often laborious to use and secondly, pre-trained models come with the danger of not being able to resolve subtle but potentially interesting heterogeneity in the data, such as a new cell state. Completely unbiased methods, such as PCA, are less liable to cause undetected heterogeneity. We hypothesize that a model zoo can solve both problems as both transfer learning and a large enough reference data set for pre-training will alleviate issues with unresolved novel heterogeneity.

Because we have many data sets per organ, we are able to use hold-one-data-set-out test splits to reflect the ability of these models to capture variance in settings with previously not seen confounding effects. Example embeddings for human and mouse lung data sets (Fig. 5a,b) show that cell types are clearly separated. We then compared reconstruction errors in cross-validation splits across commonly used model classes across 45 organs from both mice and humans, using four different (decoupled) latent space learnings. We found that linear models perform similarly to non-linear models, with median best achieved negative log likelihood across models and organs for human samples of 0.09 and for mouse samples of 0.68 (Fig. 5c,d). Best achieved negative log likelihood for human blood models was also 0.15 for linear models, consistent with the reconstruction error found on the held-out PBMC data shown in the automated example analysis above. This finding shows that single-cell data can often be represented well by linear models^13^, which have computational advantages. In sfaira, we improve this analysis in three aspects. By deploying pre-trained models that are already optimized for hyperparameters, we alleviate the need for grid searches or feature engineering. Second, we reduce the burden for model interpretation as previously annotated model components, such as bottleneck dimensions, can be easily leveraged for new analysis, thereby adding value to an analysis that goes beyond representation capabilities. Third, by enabling training on extremely diverse data sets, we pave the way for the usage of highly interpretable models that are more difficult to train. The embedding models shown here are examples of models that can be used in a model zoo but do not represent the full range of pre-trained models that could be used in the single-cell context^17^.

**Figure 5:**
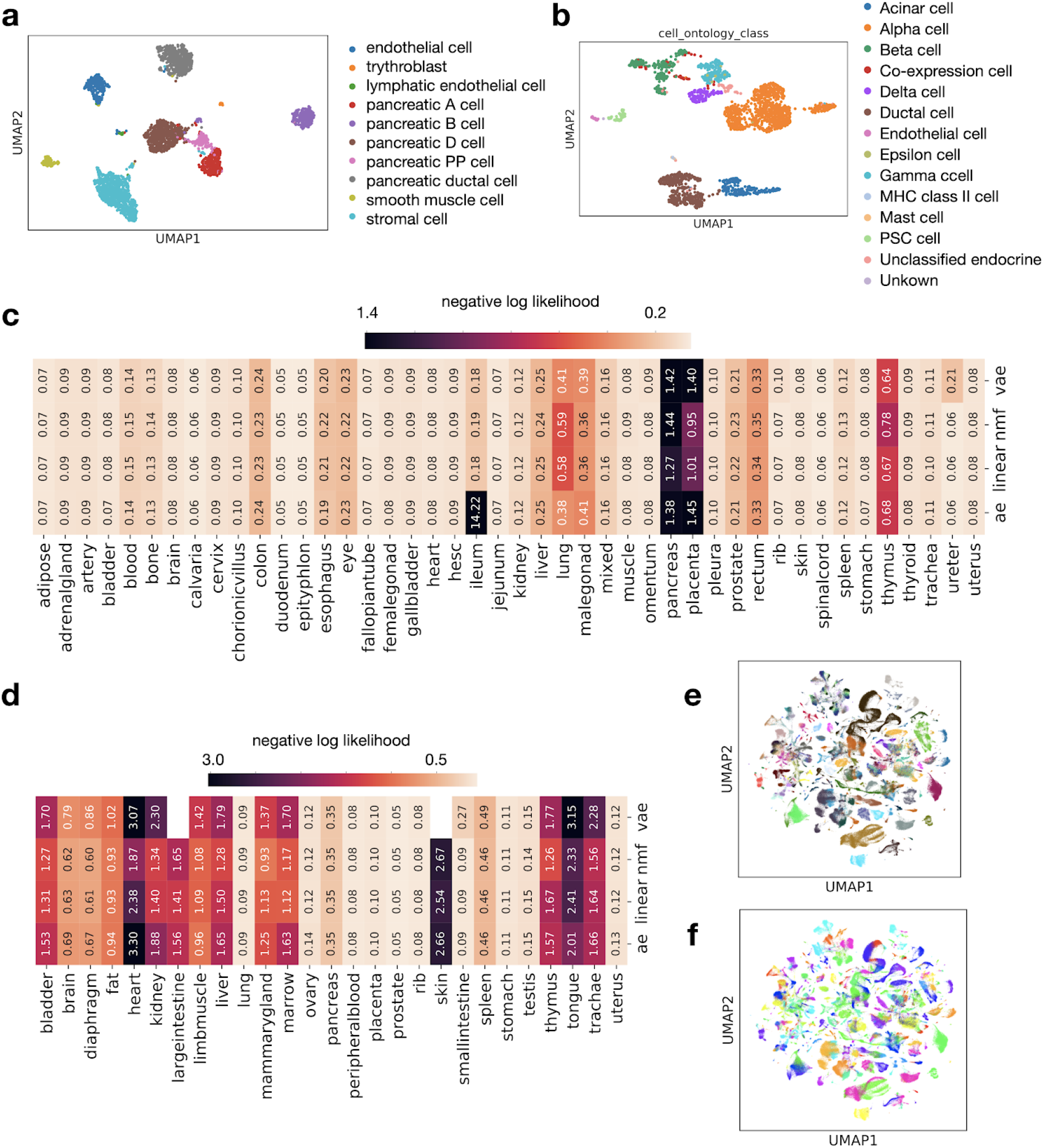
Sfaira allows streamlined embedding models training across tissues and on whole atlases. **(a, b)** UMAP on latent space using the best embedding model data for pancreas data from humans **(a)** and mice **(b).** The superimposed colors correspond to cell types. **(c, d)** Negative binomial likelihood of reconstructed test data from PCA (linear), NMF (nmf) and autoencoder (ae) models on human (**c**) and mouse (**d**) organs on entirely held-out test-data sets. **(e, f)** Sfaira allows for training of embedding models using very large data sets. Embedding model trained on all mouse data in the sfaira data zoo with the data set **(e)** and cell type **(f)** superimposed.

### Regularizing models through organism-level data

Data integration is a trade-off between removing between-sample variance resulting from technical effects and conserving biologically meaningful variance^3^. Instead of removing between-sample variance in a data integration setting, we focus on embedding models that discover axes of variation which allow us to discern biological variation on a new data set (zero-shot learning^18^). Here, it is difficult to discern models that overfit all variation in data sets and models that capture only relevant axes of variation. This overfitting is an issue that can be addressed through regularization. Model regularization in embedding and cell type classifier neural networks is often performed via L1 or L2 constraints on model parameters or via drop-out mechanisms^5^. Similarly, embedding model bottlenecks are also a form of regularization, thus limiting the effective degrees of freedom of the model to fit the data. While effective in the prevention of overfitting, these regularization methods cannot be easily used to derive interpretable models. Instead, they dynamically limit the degrees of freedom of generously over-parameterized models.

In principle, models can also be regularized through extremely diverse training data, thus making it hard for the model to overfit the entire training domain and forcing the model to learn strong abstractions. This approach stands in contrast to the usage of data set covariates, which effectively allows models to regress out between-data-set variance without attributing these differences to the gene-gene interactions that are implicitly modeled in the reference data in encoder and decoder parameters. This setting is often found in data integration studies, in which domain differences are represented as sample-level covariates and are removed by a projection mechanism^19^. We argue that to fit more interpretable models with higher levels of abstraction, one could fit condition-free models without sample-level covariates, but rather naive models, that only receive the expression state. However, the assembly of such organism-level data sets is often prohibitively labor intensive and poses practical issues in terms of data set size. These organism-level data sets can be easily assembled with the sfaira data zoo. We showed for the first time that one can converge embedding models on such data zoos of scRNA-seq data, exploiting features of sfaira and AnnData^8^ (Fig. 5e,f).

### Embedding model interpretation through gradient maps from bottleneck to input features

Many embedding models that are used in single-cell RNA-seq have been based on PCA. PCA is desirable as an embedding in terms of interpretability, because it allows for a direct interpretation of latent dimensions as orthogonal linear combinations of the input features (loadings). Gradient maps from the bottleneck activations to input features allow locally similar interpretation mechanisms in non-linear embeddings of encoder-decoder networks. Such gradient maps carry the promise of correlating bottleneck dimensions to molecular pathways or similar complex regulatory elements that present a higher-level view of gene regulatory networks. We found that cell-type wise gradient maps of the embedding space with respect to the feature space revealed cellular ontology relationships in two sample data sets (Fig. 6a, Supp. Fig. 2a) by grouping similar cell types together within a hierarchical clustering of the gradient correlation matrix. Moreover, we found that linear models and autoencoders are similar in the size of feature sets considered important by these gradient-based mechanisms for each cell type (Fig. 6a, Supp. Fig. 2a) and are also similar in the marginal distribution over normalised gradients (Fig. 6b, Supp. Fig. 2b). Models trained only with small data sets may collapse to only use small subsets of the gene space and represent cells based on feature correlations in this feature space. As data sets grow, more complex representations have to be learned, and any collapse of models on sub-feature spaces can be diagnosed with gradient-based approaches.

**Figure 6:**
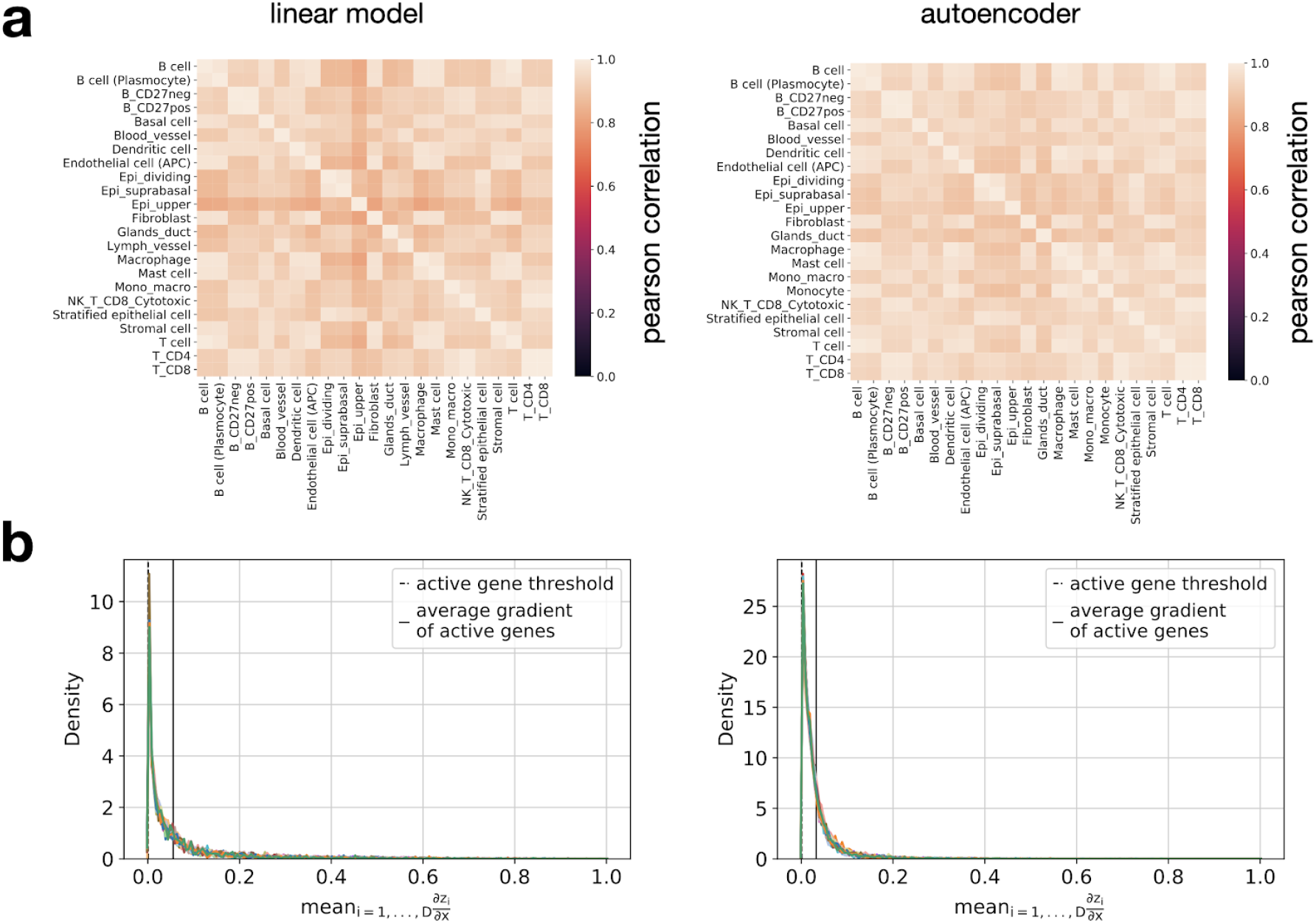
Toward the interpretability of model embedding. Saliency-based interpretation and data regularization of non-conditional embedding models: linear and autoencoder embedding models for human esophagus **(a, b)**. **(a)** Correlation of cell-type wise aggregated gradients of embedding with respect to input features. **(b)** Distribution of feature-wise wise aggregated gradients of embedding with respect to input features by cell type (color).

## Discussion and conclusions

We introduced sfaira, a data and model zoo which accelerates and standardizes data exploration for scRNA-seq data sets. The automatized exploratory analysis aspect of sfaira workflows smoothly integrates with scenario-specific scanpy^8^ workflows and scales data exploration by reducing the number of manual steps performed by analysts. Sfaira accelerates parallelized model training across organs, model benchmarking, and comparative integrative data analysis through a streamlined data access backend while improving deployment and access to pre-trained parametric models. The mechanism introduced here to accumulate large reference databases and to fit models on extremely diverse data set collections, provides a gateway to regularization through data and to mechanistic models. In contrast to query-to-reference analysis, the models presented here can be leveraged for unconstrained data exploration^19,20^. Lastly, our framework is open for the contribution of single-cell centric models that do not primarily serve the purpose of single-cell RNA-seq embedding or cell type prediction. Other use cases may include embedding models across multi-modality joint feature spaces such as CITE-seq^21^ or cell doublet prediction^21–23^.

Our effort to streamline the zoo of single-cell data is complementary to institutionalized efforts, such as the Human Cell Atlas. The sfaira data universe is currently being extended by automated data loaders that accept already streamlined data from such organizations. Our mixed data zoos can represent every data set in a publicly maintained, data-set-specific code base, and, at the same time, can leverage consistent data representations from data providers, while retaining a single interface. We expect sfaira to become a useful resource for automated data analysis, a comprehensive source of reference data sets, and to enable benchmarking of new methods.

## Supplementary Figures

**Supp. Fig. 1:**
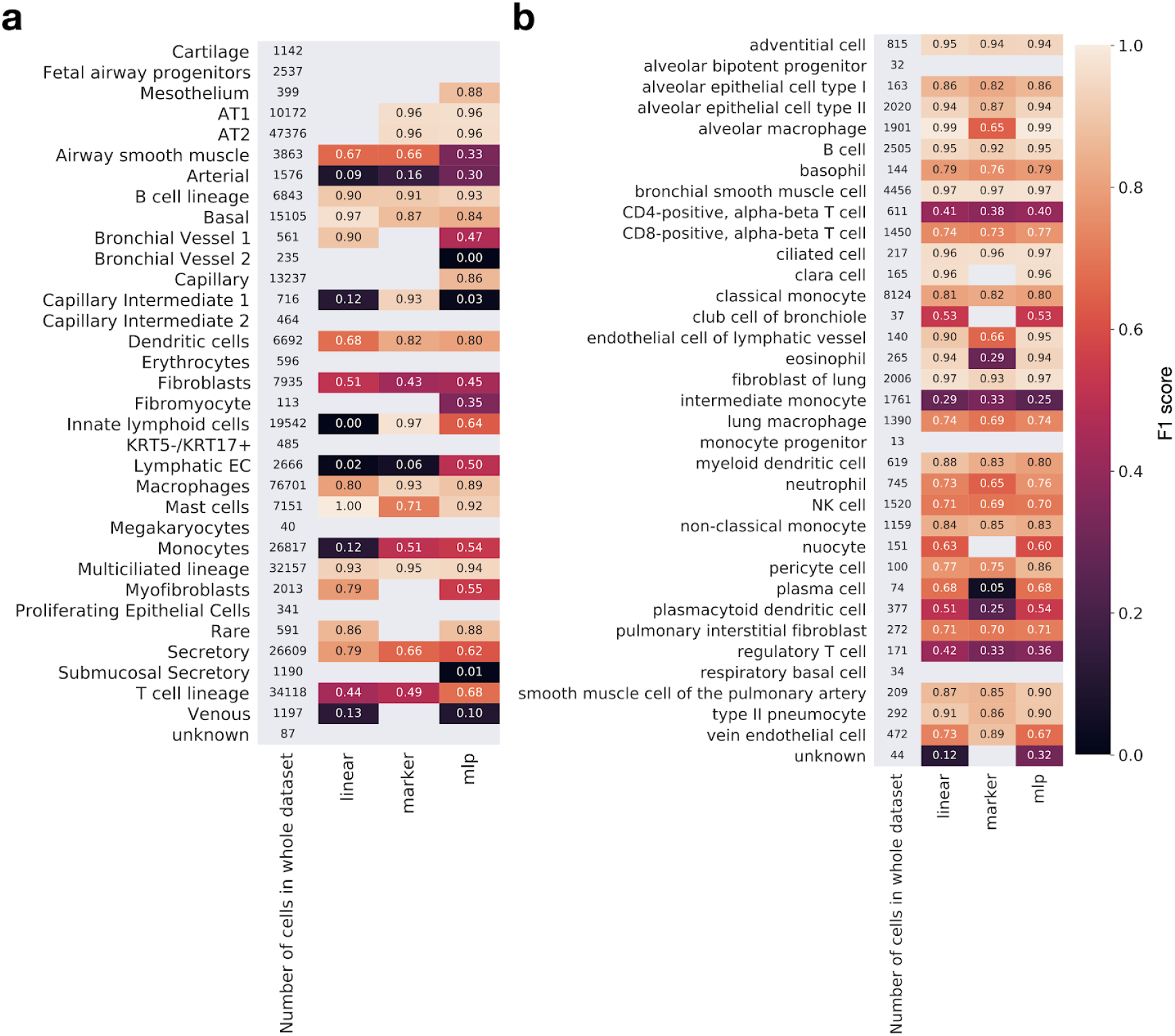
Characterization of the cell type classification task. **(a,b)** Cell type wise accuracies for lung samples from humans **(a)** and mice **(b).**

**Supp. Fig. 2:**
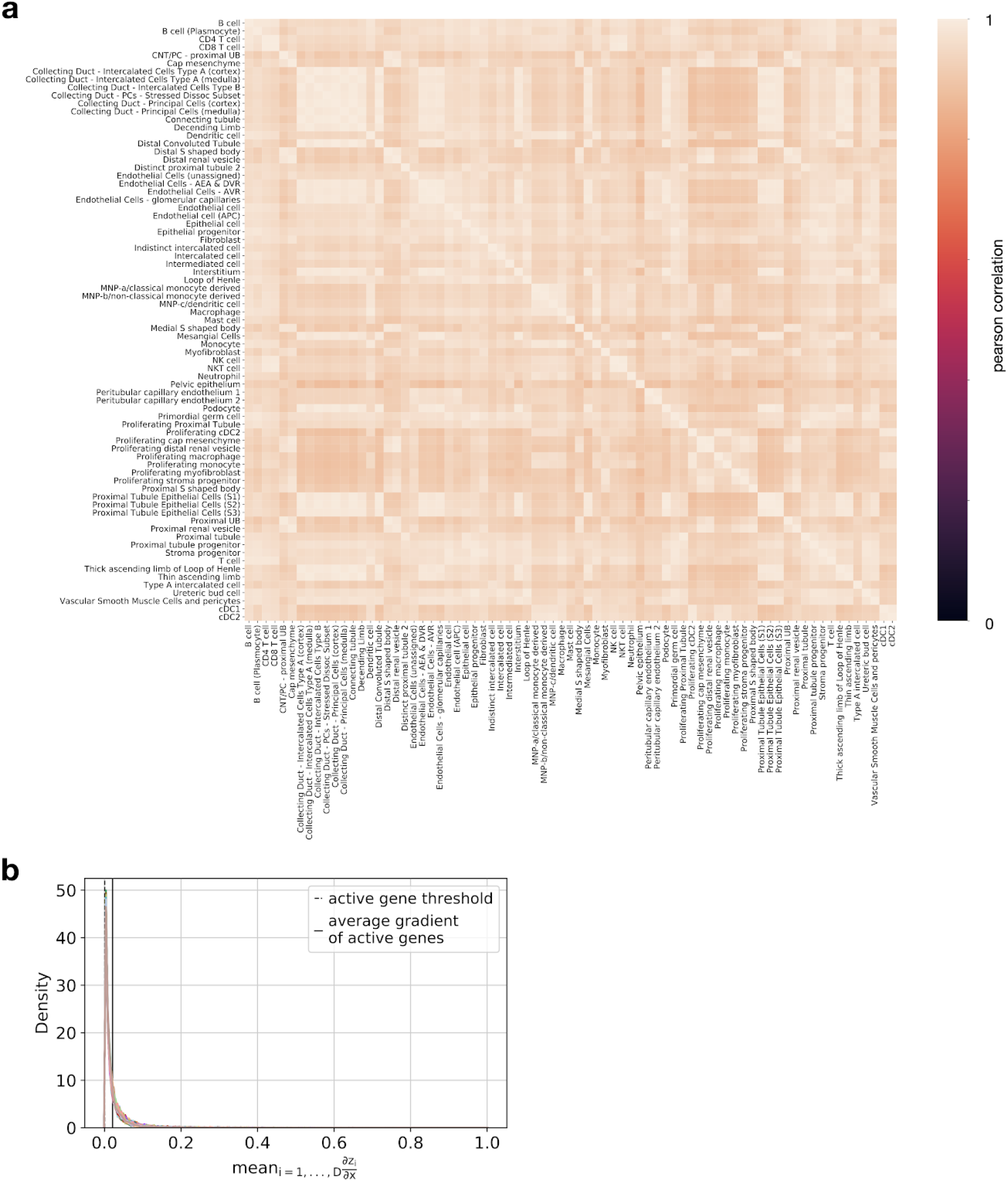
Saliency-based interpretation of models trained on human kidney. Shown are results for an autoencoder. **(a)** Correlation of cell-type wise aggregated gradients of embedding with respect to input features. **(b)** Distribution of feature-wise wise aggregated gradients of embedding with respect to input features by cell type (color).

## Declarations

### Ethics approval and consent to participate

Not applicable.

### Consent for publication

Not applicable.

### Availability of data and materials

The python package sfaira is available via GitHub (https://github.com/theislab/sfaira). An overview of the data zoo is provided here (https://theislab.github.io/sfaira-site). The notebooks containing the analysis results presented here are available at (https://github.com/theislab/sfaira_benchmarks). Data were downloaded as described in the Methods section.

### Competing interests

F.J.T. reports receiving consulting fees from Roche Diagnostics GmbH and Cellarity Inc., and ownership interest in Cellarity, Inc. and Dermagnostix.

### Funding

D.S.F. acknowledges support from a German Research Foundation (DFG) fellowship through the Graduate School of Quantitative Biosciences Munich (QBM) [GSC 1006 to D.S.F.]. D.S.F. and L.D. acknowledge support by the Joachim Herz Foundation. F.J.T. acknowledges support by the BMBF (grant# 01IS18036B, grant# 01IS18053A and grant#01ZX1711A) and the Chan Zuckerberg Initiative (grant# 2019-207271 and #2019-002438) and by the Helmholtz Association’s Initiative and Networking Fund through Helmholtz AI (grant # ZT-I-PF-5-01).

### Author’s contributions

DSF, LD and FJT conceived the study and wrote the manuscript. DSF, LD, MK, AM, LZ, ST, OH, HA wrote the python package and fitted the models.

## Acknowledgments

We would like to thank Dr. Ambrose Carr and Dr. Jonah Cool for feedback on this project.

## Methods

### Implementation

#### Data zoo

We represent data sets by individual data loader classes that inherit generic data-loading properties from a unique parent class. These data loader classes can be considered class versions of data-loading scripts that are otherwise often used in script code. These classes allow metadata queries through automatic metadata storage in a lazy mode, in which count matrices are not yet loaded into memory, thereby allowing the user to subset large instance lists of these classes interactively. Some entities serve streamlined processed data sets for which individual loading scripts are not necessary: In these cases, we interface these data zoos with a single class that can be instantiated for all data sets in this zoo. Sfaira maintains the universe of all contributed data loader classes; users then locally build libraries of a subset of these data sets, and the sfaira data api accesses all available data sets: This allows users to also only operate on a subset of the available processed data universe.

#### Model zoo

We provide a model code in the sfaira package; each model has its own model class that can be accessed through a streamlined interface, such as in *kipoi^7^*.

#### Parameter storage

Parameter files of models defined in the sfaira model zoo are stored in public cloud servers, such as Zenodo, or locally for private models. These parameter files are versioned and can thus be reproducibly accessed.

### Model topologies

Sfaira is a model zoo that is set up to accommodate various models. Here, we describe the models that underlie the analysis results that are presented in this manuscript. Note that the models in sfaira will not be limited to this initial model population in the future. One can, however, query those exact models at any point in the future by using the unique model identifiers supplied in Supp. Data 1.

#### Preprocessing layers

We prepended a common input data transform to all embeddings and cell type prediction models. The objective for using this transform is to reduce variability in the data so that models require lower complexity and fewer training steps to adjust their internal normalization of the data. We chose a transform that can be evaluated based on a data batch without being dependent on the batch. For arbitrary batch sizes, this requires the transformation of an individual observation (cell) to only depend only on the observation itself. We linearly scaled the data points *x* per cell *i* and gene *j* to 10,000 and log transformed this scaled vector.

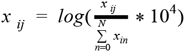

The scaling is a basic attempt to reduce the variation caused by the number of UMIs observed per cell which depends on technical factors such as the library depth and stochasticity in mRNA capture during the sequencing experiment. The log transform is a basic attempt at reducing the strong heteroscedasticity of the data which is commonly observed to have a positive dependence of the variance on the mean of the gene observations.

We would like to highlight that unlike in standard single-cell RNA-seq data processing for PCA and downstream t-SNE or UMAP computation, this processing does not necessarily need to be carefully benchmarked as this processing is complemented by the innate ability of the first layers of the neural network to adjust to unwanted sources of variation. We chose to use a basic transform to speed up training only. In the limit of many data sets and sufficient training time, one could imagine to entirely remove preprocessing from these networks.

#### Output and loss function of embedding models

We provide support for different model outputs and loss functions. These variations are encoded in the topology identifier. Multiple studies have found that autoencoder can learn embeddings of single-cell RNA-seq data with negative binomial reconstruction loss. A negative binomial reconstruction loss requires a mean and a dispersion parameter to be estimated. In the initial version of sfaira, we support output states tailored to the negative binomial distribution through an exponential inverse-linker function in the last layer. We distinguish an output that estimates a fixed dispersion per gene and an output that estimates one dispersion parameter per gene and cell.

#### Output and loss function of cell type prediction models

The standard cell type prediction model included in sfaira operates under the assumption that a cell type prediction should output a probability distribution across previously known cell types and an “unknown” or “not-a-cell” class. Such a prediction can be achieved with a softmax transform of the output states in the layer, the loss can be evaluated based on cross-entropy loss. We additionally allow for multiple output categories to be assigned to a single true set of labels, we call this aggregated cross-entropy loss and we use aggregated accuracy as an evaluation metric for this scenario. This aggregation is necessary if data sets differ in the coarseness of the cell type assignments. Often, one can map labels between both data sets as part of an ontology. Data set A may only annotate four tissue-specific cell types and “lymphocytes” whereas data set B differentiates those four types and further differentiates “T-cells” and “B-cells”. The cell universe of this tissue should, therefore, consist of the four tissue specific cell types and T-cells and B-cells. Data set A can still be used to train supervised classifiers to predict cell types, but one must take care that the lymphocyte label is used properly. We propose to aggregate the predicted class probabilities across all labels assigned to lymphocytes in data set A so that any probability mass distribution for a lymphocyte observation in A across T-cells and B-cells is allowed. This allows the classifier to learn differences between T-cells and B-cells on data set B, while it can use A to improve its model of the difference between both lymphocytes and the remaining four cell types. Below, we compare the resulting aggregated cross-entropy loss *cce*_*agg*_ to cross entropy for a binary (*cce*_*binary*_) and a multi-class (*cce*_*multi*–*class*_) prediction problem. The shown transformations labeled with (*) hold if *y* ⊂ {0, 1}, ie., if the labels lie on a binary support.

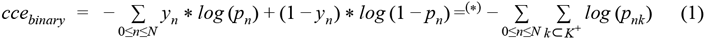

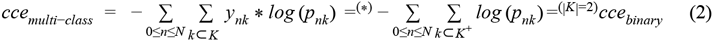

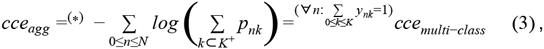

where *K*^+^ is the set of positive classes with *y*_*k*_ = 1 and *K*^−^ is the set of positive classes with *y*_*k*_ = 0 and *N* is the number of observations. In the above example, 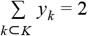 for observations assigned as “lymphocyte” for both the T-cell and B-cell entry in *y* would be equal to one. Observations assigned as “T-cell” would be calculated as 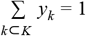. The accuracy metric *acc*_*agg*_ corresponding to *cce*_*agg*_ is:

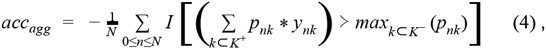

where *I* [] is the indicator function.

Alternatively, one could use sigmoid transforms of independent cell type predictions. This approach does not superimpose the prior knowledge that a cell can only be part of one class in a properly defined cell type ontology, and thus, we therefore do not support this setting.

#### Autoencoders

Autoencoders with “dense” (fully-connected) layers and count noise distributions were proposed among others by Esralan *et al.* to learn embeddings of single-cell RNA-seq data^5^. The full architectures are documented in the code.

#### Variational autoencoders

Variational autoencoders (VAEs) with “dense” (fully-connected) layers on count noise data were proposed among others by Lopez *et al.* to learn embeddings of single-cell RNA-seq data^6^. Here, we impose a unit Gaussian prior on the latent space activations. The full architectures are documented in the code.

### Data processing

#### Expression data

All data (human^24–56^ and mouse^57–61^) were downloaded in the least-processed expression matrix format provided by the authors of the data set. We did not perform any processing other than that discussed for preprocessing layers discussed in the section “Model topologies”. As feature space we chose the protein coding genes from the GRCm38 genome assembly for mice and GRCh38 for humans.

#### Cell type annotation data

Not all data sets used in this manuscript use the same cell type identified conventions. We manually assembled a baseline set of cell types per species and tissues from the entire input data and mapped all individual labels to this reference set using the data reading functions that we provide in the Github repository. In some cases, the provided annotation only contains superset identifiers, such as lymphocytes for an organ for which we have T-cell, B-cell, and macrophage labels. In these cases, we assigned a positive label of 1 to the set of labels that maps to this superset label (see also aggregated cross-entropy, Methods eq. 3).

